# Association Between Oral Microbiota and Close-Range Proximity in a Primary School

**DOI:** 10.1101/2024.12.27.628096

**Authors:** Lorenzo Dall’Amico, Xiangning Bai, Sandra Marie Weltzien, Simon Rayner, Daniela Paolotti, Isabelle Sylvie Budin Ljøsne, Andreas Matussek, Anne-Sofie Furberg, Ciro Cattuto, Christopher Sivert Nielsen

**Affiliations:** ISI Foundation, Turin, Italy; Department of Microbiology, Division of Laboratory Medicine, Oslo University Hospital, Oslo, Norway; Division of Infectious Diseases, Department of Medicine Huddinge, Karolinska Institutet, Stockholm, Sweden; Faculty of Health Sciences and Social Care, Molde University College, Molde, Norway; Department of Medical Genetics, Oslo University Hospital and University of Oslo, Oslo, Norway; Institute of Clinical Medicine, University of Oslo, Oslo, Norway; Department of Food Safety, Norwegian Institute of Public Health, Oslo, Norway; Department of Microbiology, Division of Laboratory Medicine, Institute of Clinical Medicine, University of Oslo, Oslo, Norway; Department of Microbiology and Infection Control, University Hospital of North Norway, Tromsø, Norway; Department of Chronic Diseases, Norwegian Institute of Public Health, Oslo, Norway; Department of Pain Management and Research, Oslo University Hospital, Oslo, Norway

## Abstract

The microbiota is the ensemble of microorganisms inhabiting the human body and it deeply influences human health and well-being. Recent studies showed its interplay with social behavior, suggesting that part of the microbiota might be socially transmissible. In this work, we investigate the association between close-range proximity and the oral microbiota composition in a group of children attending primary school. Unlike most related studies, our cohort comprises non-cohabiting individuals, and we use high-resolution proximity sensors to objectively quantify close-range proximity. Our analysis shows that longer time spent in proximity between pairs of children correlates with a higher similarity between their oral microbiota composition. Building on previous studies on cohabiting individuals, our findings suggest that social contacts beyond the closest social circle may also shape oral microbiota composition, although longitudinal studies are needed to test this hypothesis.

**Author summary:** The human microbiota is the ensemble of microorganisms inhabiting the human body. It significantly influences human health and understanding the determinants of its composition is consequently a relevant open question. Dietary and environmental factors are known to play a key role, but recent evidence emphasized the role played by social interactions. In isolated communities, strong bonds, such as familiar bonds, are predictors of microbiota formation. In our work, we consider a highly non-isolated social environment, composed of non-cohabiting children in a primary school. We show that the oral microbiota composition predicts close-range proximity relations. Our results evidence the strong relationship between close-range proximity and human microbiota composition for social contacts reaching beyond the closest social circle.

## Introduction

The human microbiota is a complex ecosystem of microorganisms inhabiting the human body. It is predominantly composed of bacteria but also includes viruses, archaea, and fungi that engage in a continuous interplay of cooperation and competition [1]. It is known that the human microbiota significantly influences the host’s phenotype, energy metabolism, immunity, and, ultimately, human health and well-being [2, 3, 4]. Recent studies also demonstrated its impact on psychological development and social behaviors in humans and animals [5, 6, 7, 8, 9]. As such, the human microbiome is considered the second human genome [10].

Social interactions and human health are deeply intertwined and are known to influence one another. Seminal studies have shown that obesity [11] and depression [12] are shaped by social interactions and can spread in a community. Likewise, a growing body of literature suggests that social interactions can influence the composition of human microbiota [13]. In humans, microbiota is exchanged both vertically – primarily from the mother – and horizontally through interactions with peers [14]. Over time, horizontal exchange becomes the dominant process, and differences in dietary habits, environmental factors, and social contexts lead to differences in the microbiota between the child and the parents [15].

The association between microbiota composition and social structure was first documented in several non-human mammals, including mice [16], baboons [17, 18], chimpanzees [19], and lemurs [20], among others. To date, only a handful of studies have investigated the correlations between human microbiota and close-range social interactions. Ref. [21] showed that the skin microbiota of people moving to a new house rapidly aligns with that of the previous tenants. Similarly, Ref. [22] observed that family members and their dogs living in the same household share skin microbiota [22]. Other studies showed that spouses can be identified from their skin microbiota [23], and that they have more similar gut microbiota and share more bacterial taxa than siblings [24]. These works suggest that, in adult life, prolonged interactions and similar lifestyles play a predominant role in shaping microbiota.

In the case of non-cohabiting individuals, Ref. [25] investigated the relationship between the oral microbiota and the social network in a group of hunters and gatherers. They introduced and identified the *social microbiome* as a portion of the oral microbiome hypothesized to be transmitted through social interactions. The authors also observed a higher diversity in the oral microbiota composition of the most central nodes in the social network, in agreement with Ref. [8], which obtained a similar result for the gut microbiota.

The cited works reported an association between the microbiota composition and the social structure, while only in a few cases transmission could be assessed [26, 27]. Ref. [26] showed that it is possible to infer cohabiting couples from their gut microbiota, while Ref. [27] found greater gut microbiota similarity among individuals having frequent interactions in a group of isolated villages.

Although the correlation between social interactions and the microbiota composition has been observed in many studies, research in this field is still in its early stages because of the many technical challenges it poses. One challenge is the accurate measurement of close-range proximity relations. In previous studies, social interactions were typically defined in terms of known, long-standing relations or proxies thereof. The most frequent contacts were usually identified by direct observation [18, 19] or using surveys, contact diaries, and self-reported information on interpersonal affinity between individuals [27]. These measurements are subjective and tend to be biased towards long-lasting relations [28, 29]. In many cases [15, 16, 20, 22, 26, 24], membership in a family/household/village was used as a proxy for interaction. While information on these associations is readily available and reliable, using such strong relationships does not allow the study of the microbiota’s similarities (or lack thereof) among individuals who have social contacts reaching beyond their closest social circle. One exception is Ref. [25] in which proximity contacts were objectively measured with radio sensing technology.

In this work, we investigate the relationship between social interactions and oral microbiota composition in a group of children attending primary school in Molde, Norway. We focus on close-range proximity happening at school, and we measure them using the high-resolution proximity sensors developed by the `SocioPatterns` collaboration [30, 31]. We analyze the oral rather than the gut microbiota because changes in its composition are more likely to be observed in relatively short time frames [15]. Moreover, based on long-term studies of individuals in a confined environment [32], it was suggested that social influence might impact the oral microbiome more than the gut microbiome. It is also worth mentioning that, although oral microbiota appears to be more easily transmitted, a strong relationship has been shown between oral and gut microbiota [33].

## Results

### Data description

Our data describe a group of 35 children from two third-grade classes at Nordbyen primary school, Molde, Mid-Norway. During class hours, the children sat in pairs in fixed positions that changed every two weeks, except for group work and practical teaching sessions. They engaged in outdoor and indoor free activities during breaks and in the after-school care program. This cohort comprises individuals from different families, allowing us to investigate the correlation between close-range social contacts and oral microbiota among people who share a common environment, a school class in our case. Additional details on the cohort composition and the enrollment procedure can be found in the Methods section.

We characterized participants’ oral microbiota through saliva samples collected during class hours at eight time points. Due to data quality issues, we consider only the first five samples, collected between September 12 and October 7, 2022. Microbial composition was profiled using 16S rRNA gene sequencing, yielding relative abundance distributions for each taxon [34], recorded at the amplicon sequence variant (ASV) resolution. Across all samples, we identified 12,322 amplicon ASVs.

We compare microbiota samples using three metrics: the Jaccard distance between the sets of all taxonomic labels present in two samples; the Bray-Curtis dissimilarity [35], computed on relative abundances; and the weighted UniFrac distance [36], which incorporates the phylogenetic relationships between taxa.

We assess microbiota stability over time and across individuals. Fig. 1A shows that each individual’s microbiota composition remained remarkably stable throughout the study period, across all three metrics. This is consistent with previous evidence on similar timescales [15, 37]: samples from the same individual show significantly lower distances than those from different individuals (Mann–Whitney test; rank-biserial correlation of 0.91 for Jaccard, 0.84 for Bray-Curtis and 0.53 for weighted UniFrac, with *p*-values below machine precision). We reinforce these results with PERMANOVA analysis. The variation between samples is well explained when grouping by individual identity (*R*^2^ = 0.55 for Jaccard, *R*^2^ = 0.69 for Bray-Curtis, *R*^2^ = 0.55 for weighted UniFrac with *p*-values ≤ 0.001), but not when grouping by sampling date. Only weighted UniFrac shows a significant effect of sampling day, with a substantially smaller effect size than that of individual identity (*R*^2^ *≈* 0.04, *p*-value *≈* 0.01). In the Supplementary information we show the robustness of our results when filtering rare variants, which likely arise from sequencing errors or contamination [38], and non-colonizing variants. We remark that the unweighted UniFrac distance is not considered because, in our data, it is not robust when applying these filters, unlike the other three metrics. In the Supplementary information, we further demonstrate, using different alpha-diversity metrics, at every filtering level, that there is no significant difference in microbial diversity across measurements.

**Figure 1:**
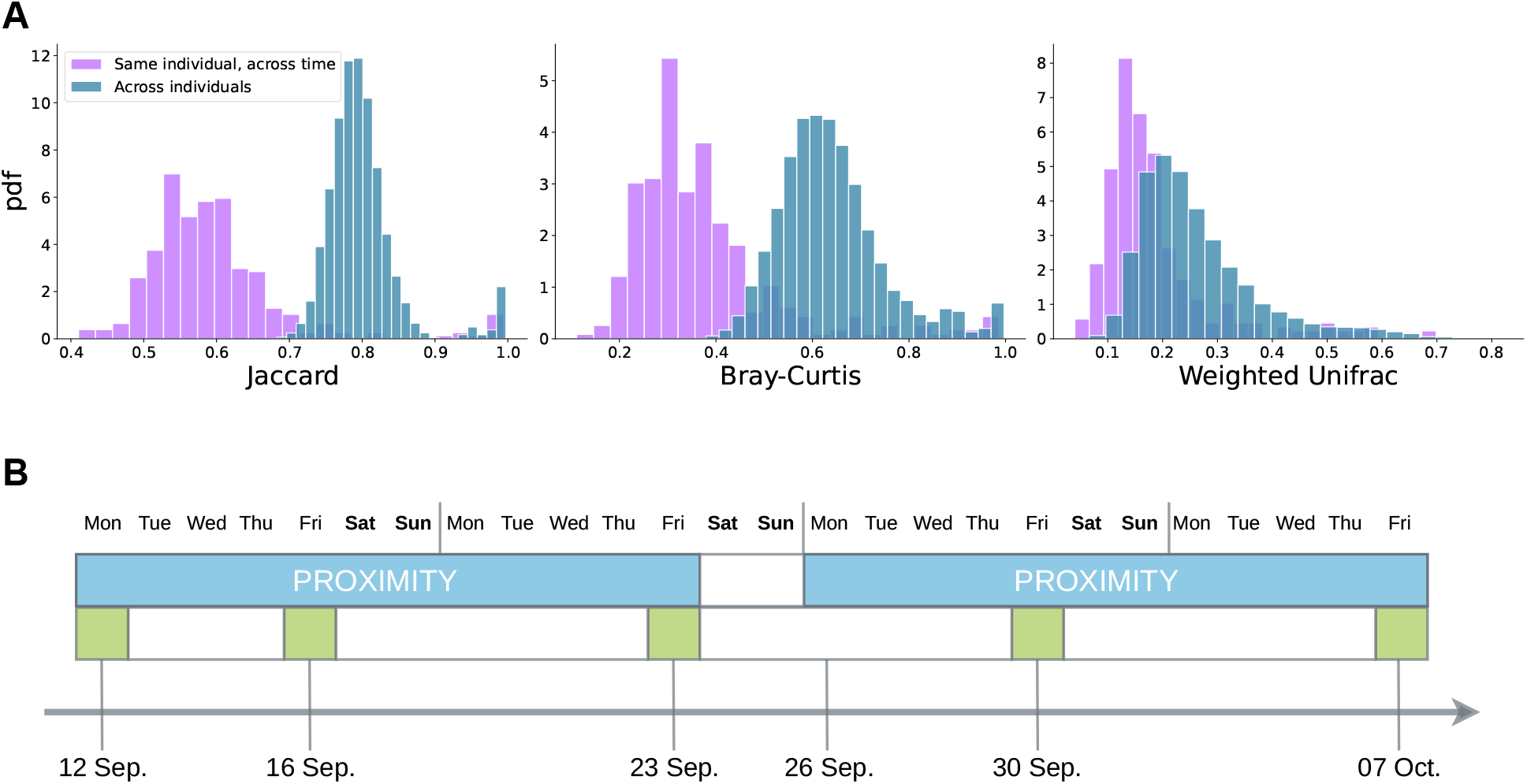
Microbiota stability and data collection timeline: ***Panel A***: *Histograms of microbiota distance across individuals and across time for the same individual, for three distance metrics*. For each of the Jaccard, Bray-Curtis, and weighted UniFrac distances, the histograms represent the distance between all measurements involving the same person (“Same individual, across time”, purple) and between all measurements involving different individuals (“Across individuals”, blue). ***Panel B***: *Data collection timeline relevant for this study*. The arrow at the bottom indicates time and reports the dates in 2022 relevant for data collection. The light blue bars indicate the time intervals during which proximity sensors were used. Saliva samples were collected on the days highlighted in green, on the bottom row.

In parallel with the microbiota sampling, we measured a short-range proximity network among children with close-range proximity sensors, developed by the `SocioPatterns` collaboration [30, 31]. These sensors are a well-established, unobtrusive, scalable technology that has been used to quantify human close-range proximity in a variety of real-world settings spanning schools [39, 40, 41], offices [42, 43], hospitals [44] and households in low-resource settings [45, 46, 47, 48], among others. Like the microbiota sampling, the proximity measurements were collected in 3 waves. Due to accumulating damage to proximity sensors (see Methods section), only the first two waves achieved satisfactory coverage of the participants’ contacts. The total time in proximity across the two measurements is highly correlated (Spearman *r* = 0.25 with *p* value 7*·* 10^*−*14^). Therefore, we choose to work with static representation of contacts obtained by aggregating all the proximity information collected by the proximity sensors into a single, undirected, weighted social graph. Graph nodes represent children, edges are proximity relations, and edge weights are defined as the total time spent in close-range proximity. Reported proximity durations are restricted to school hours: off-school contacts were measured but contributed with a negligible fraction of the total contacts, hence we excluded them from our analysis. We focus on the five microbiota sampling times that fall within the period covered by the first two waves of proximity sensor wearing and, given the stability of the microbiota measurements (Fig. 1), we quantify microbiota similarity by averaging the similarity across the five samples. Fig. 1B provides an overview of the data collection schedule used in this study.

### Microbiota similarity as a function of close-range proximity

We want to study the relationship between microbiota composition and close-range proximity, testing the hypothesis that the microbiota composition is more similar for pairs of subjects who spend a longer time in proximity. To address this, we construct two social graphs encoding, respectively, physical proximity and microbiota similarity. Nodes represent the students, and both graphs are complete, *i*.*e*., connections exist between all pairs of nodes. The two graphs differ in how edge weights are defined. In the microbiota similarity graph, denoted by *G*_m_, the edge weights are defined as the average distance (Jaccard, Bray-Curtis, or weighted UniFrac) across the five samples between the microbiota compositions of the incident nodes. In the proximity graph, denoted by *G*_p_, edge weights increase with the total time in proximity expressed in seconds, *τ*, between the incident nodes. It is well known that in human proximity networks, the probability distribution of contact durations *τ* is fat-tailed [31, 39, 41, 42, 45, 47], and we consequently use a logarithmic weighting for the edges of *G*_p_, expressed as *w* = log(1 + *τ*). This expression leads to zero edge weights for subject pairs that did not come into proximity according to the sensors (*τ* = 0).

To test whether pairs with longer contact duration have more similar microbiota compositons, we focus on the strong links of the proximity graph, *i*.*e*., on edges corresponding to the longest-lasting proximity relations, as measured by the wearable sensors. We introduce a threshold proximity duration, *τ*\*, and define “strong” edges as those with duration *τ > τ*\*. We filter the proximity graph according to this threshold, retaining only strong edges, and study the average microbiota distance among the reained edges as a function of the threshold *τ*\*, for each of the three distance metrics. To assess the significance of our findings, we compare the observed values with a null model that randomizes the association between the weights of *G*_p_ (related to contact duration) and *G*_m_ (microbiota distance), over 1500 realizations. Fig. 2 shows that, for all three metrics, the average microbiota distance among strong ties decreases with increasing duration threshold *τ*\*, and falls outside the null model’s confidence band for intermediate values of *τ*\*, approximately between 2 and 4 hours. We remark that for large values of the duration threshold, *τ*\* *≥* 5 hours, the results are only bordeline significant or non-significant. This result is expected, as for such large values of the threshold, only a very small portion of the graph’s edges (approximately 3%) are retained for the analysis, and noise tends to dominate the results. The Supplementary information shows that these results are robust when excluding rare or non-colonizing variants.

**Figure 2:**
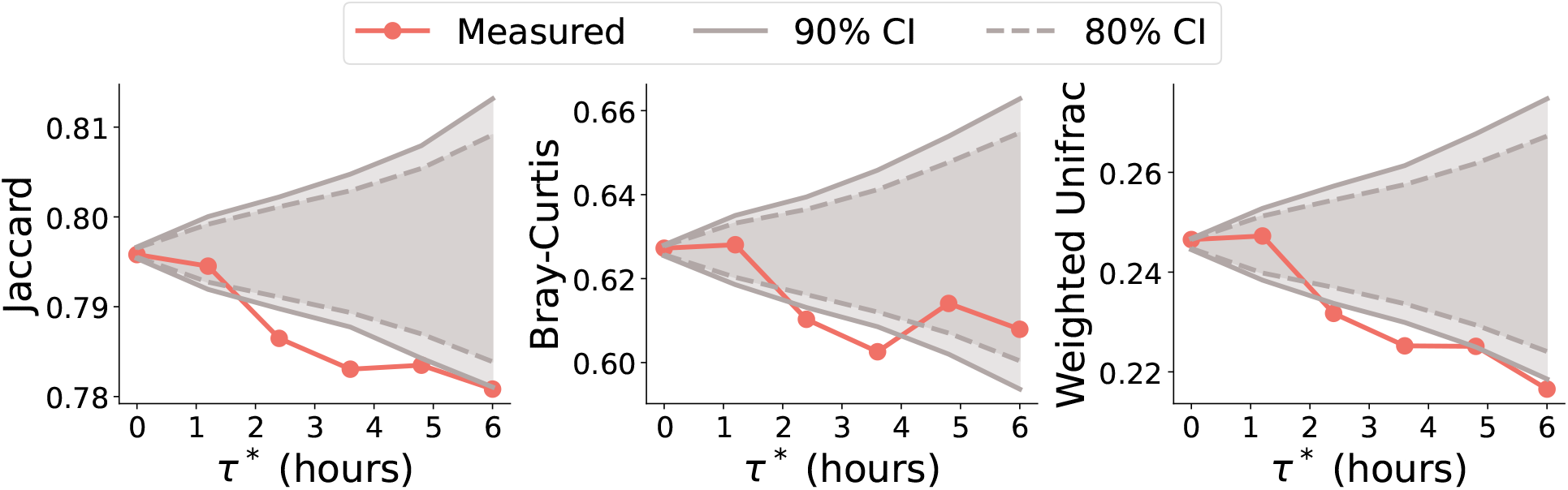
Microbiota similarity between node pairs forming strong ties, as a function of the definition of strong tie, for three distance metrics. ***Panels A–C***: *Jaccard, Bray-Curtis, and weighted UniFrac, respectively*. Strong ties have a duration *τ* larger than the threshold *τ*\*, spanning the *x*-axis. The *y*-axis shows the average microbiota distance (weight of *G*_m_) between nodes forming strong edges, for the corresponding metric. The two style-coded gray lines are the 80% and 90% confidence intervals obtained over 1500 realizations of the null model randomizing the association between the weights of *G*_p_ and *G*_m_. Proximity strength increases from left to right, whereas microbiota similarity increases from top to bottom.

### Inference of the proximity network from the microbiota

We further explore the relationship between oral microbiota similarity and close-range proximity by studying whether we can predict the proximity network using the oral microbiota information alone. Specifically, we train a binary classifier that predicts, given the microbiota profiles of two individuals *i* and *j* and a duration threshold *τ*\*, whether a link between *i* and *j* with weight (duration) exceeding *τ*\* is present in the measured proximity network. Missing links in the network are treated as zero weight edges. In other words, given a threshold duration *τ*\*, we consider the complete graph of relations between individuals, and assign a link (*ij*) to class “1” if nodes are *i* and *j* were observed in proximity for a total duration longer than *τ*\*, and to class “0” otherwise. As an input to the classifier, we use a binary feature vector ***y*** describing the taxa sharing between nodes *i* and *j*: entry *y*_*a*_ of ***y*** is 1 if taxon *a* was hosted at any time by both *i* and *j*, and 0 otherwise. To deal with the high dimensionality of ASV data, we adopt the standard machine learning approach of using a support vector machine (SVM) [49] trained on a reduced feature space comprising the 50 largest principal components of the matrix formed by the vectors { ***y***}. We validate the performance of the classifier by cross-validation, training it on a random sample of 90% of the links and testing it on the remaining 10%. Fig. 3 shows the true positive rate (TPR) versus the false positive rate (FPR) for 100 random realizations of the training and test sets, for varying values of the threshold *τ*\*. The classifier achieves an average area under the curve (AUC) of 0.77. We remark that, while the accuracy of the inference task generally depends on the algorithm adopted, a single instance in which a classifier can be successfully trained (in our case, the SVM) is enough to assess the inference feasibility, hence to show the association between proximity and the microbiota composition. The Supplementary information shows the ROC curve for a random forest classifier, which obtains a similar performance (AUC = 0.74).

**Figure 3:**
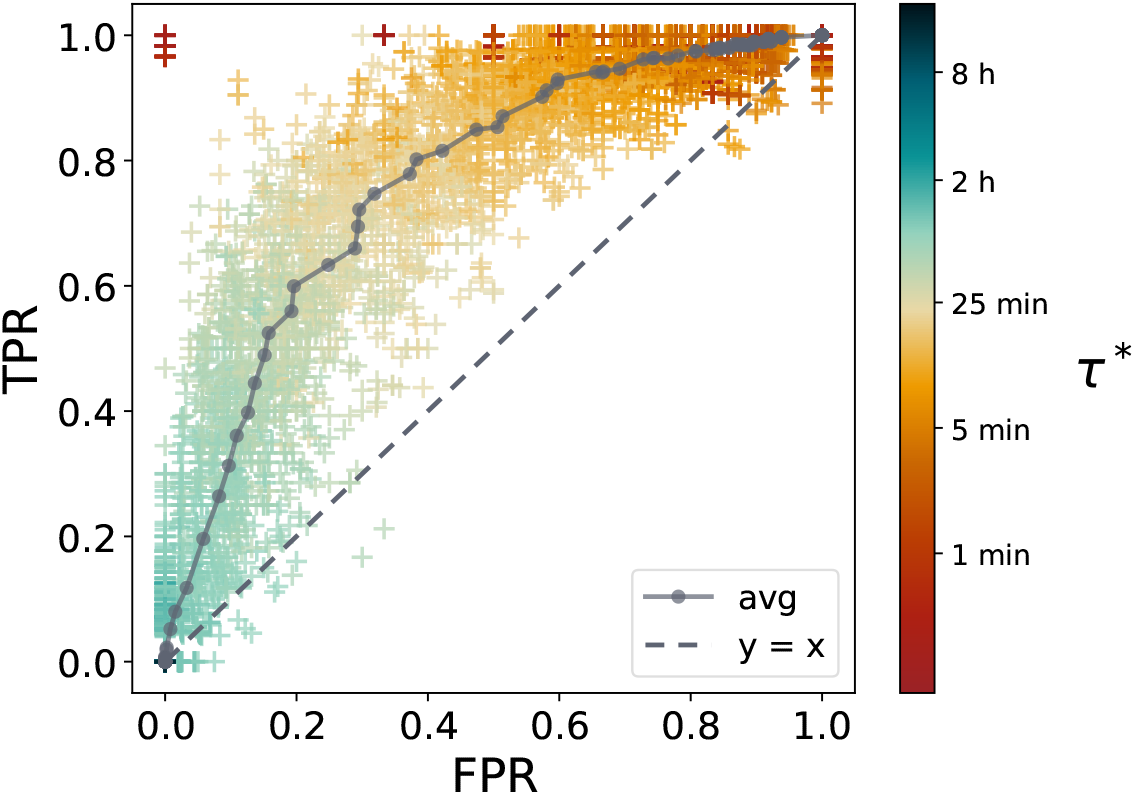
ROC curve for the binary classifier used to infer the proximity links based on microbiota profiles. We train a support vector machine to predict whether the weight (duration in proximity) of a given edge (*ij*) exceeds a threshold *τ*\*. We use, as feature vectors, the first 50 principal components of the one-hot encoding vectors indicating the ASVs shared by node pairs. The classifier is validated by training on a random selection of 90% of the links and tested on the remaining 10%. 100 random realizations of the training/test split are used, for each of the 250 logarithmically spaced values of *τ*\* we scan. Every cross in the plot corresponds to one realization of the training and test sets. The *x* coordinate is the false positive rate (FPR) and the *y* coordinate is the true positive rate (TPR). Markers are color-coded according to the value of the threshold *τ*\*. The solid gray line represents the average over the random splits and has an area under the curve auc = 0.77. The dashed line is the curve *y* = *x* for a random-guess binary classifier.

### Identification of taxa more strongly associated with network structure

In this section, we identify a set of taxa whose co-occurrence in pairs of individuals correlates with sustained close-range physical proximity. We independently consider each variant *a* and compare the proximity duration of edges in which both nodes hosted that variant (denoted with *y*_*a*_ = 1, following the notation of Section **Inference of the proximity network from the microbiota**) with those of edges in which only one of the two nodes hosted that variant (denoted with *y*_*a*_ = 0). We test the hypothesis that the distribution of microbiota variants is homophilic over the proximity network, *i*.*e*. that stronger proximity is associated with similar attributes. We thus identify the taxa *a* more strongly associated with social contacts as those in which the *y*_*a*_ = 1 edges have a significantly longer duration than those with the *y*_*a*_ = 0 edges. We attribute a duration equal to zero to the edges that were not measured by the sensors. Note that we exclude all edges in which neither of the two nodes ever hosted the considered variant, since such pairs carry no information about whether variant sharing is associated with contact duration. We employ the Mann-Whitney signed test to assess the significance of the difference between the two distributions. This is a non-parametric signed test allowing us to test if the edges in which the nodes share a taxon (*y*_*a*_ = 1) have *longer* durations than the others. We repeat the test independently on each taxon, and apply the Bonferroni correction to the statistical significance of the Mann-Whitney test. The test shows that edges with *y*_*a*_ = 1 (those in which both nodes hosted the variant *a*) have longer durations for several taxa. Table 1 reports the last available taxonomic level for all the taxa in which the *p* value is smaller than 0.05, after the Bonferroni correction, the corresponding rank bi-serial correlation, and the size effect express as difference between the medians of the two distributions. For each variant, we also report the fraction of carriers in whom the taxon was detected persistently (in at least 3 of the 5 samples) rather than only occasionally. Nine of the twelve taxa are majority-persistent (53–86% of carriers), two are borderline (40–43%), and one is persistent in only about 11% of its carriers. The prevalence ranges between 46% and 95%. The Supplementary information further shows that the identity of the taxa shared between repeated measurements from the same individual varies moderately (median Jaccard index ranging between 0.54 and 0.60), but their similarity does not decrease over time. This suggests that there is no sizeable taxon turnover over the timescale of our study. Fig. 4A compares the duration distributions for the edges with *y*_*a*_ = 1 and *y*_*a*_ = 0 for the variants reported in Table 1, and Figure 4B provides a visualization of the persistence of each variant in the population. These results show an association between the microbiota and prolonged close-range physical proximity. We note that some of the genera in Table 1 were also identified as part of the “social microbiome” described by [25], where they were associated with periodontal diseases. This is particularly the case of *Corynebacterium, Capnocytophaga, Porphyromonas*, and *Prevotella*. We remark that we identified three variants (*Prevotella pallens, Prevotella melaninogenica, Clostridiales bacterium*) for which the Mann-Whitney is significant under the hypothesis that *y*_*a*_ = 1 edges have *shorter* durations.

**Table 1:** List of taxa more strongly associated with the proximity network. The column “Taxon” is the variant identifier, which is preceded by a letter denoting the last available taxonomic level. The letter “*g*” stands for genus, and “*s*” for species. The second column reports the *p* value of the Mann-Whitney test after the Bonferroni correction, while “Corr.” is the corresponding rank bi-serial correlation. The column “*δ*_*t*_” reports the difference between the medians of the two distributions, expressed in minutes. The column “Persistence frac.” reports the fraction of the population for which that taxon was measured at least 3 out of 5 times. Values above 0.5 are underlined. The column “Prevalence” indicates the fraction of individuals in which the taxon was detected at least once.

| level | taxon | $p$ value | Corr. | $\delta_t$ | Persistence frac. | Prevalence |
| --- | --- | --- | --- | --- | --- | --- |
| g | Streptococcus | 7.81E-13 | 0.60 | 25 | 0.43 | 0.95 |
| s | Veillonella | 6.92E-08 | 0.35 | 22 | <u>0.66</u> | 0.86 |
| g | Porphyromonas | 4.46E-06 | 0.29 | 23 | <u>0.54</u> | 0.70 |
| s | Selenomonas | 4.53E-05 | 0.27 | 20 | 0.40 | 0.68 |
| g | Porphyromonas | 1.08E-03 | 0.24 | 18 | <u>0.68</u> | 0.76 |
| s | Corynebacterium | 5.67E-03 | 0.23 | 16 | <u>0.58</u> | 0.84 |
| s | Capnocytophaga | 8.93E-03 | 0.22 | 16 | <u>0.62</u> | 0.78 |
| g | Streptococcus | 1.19E-02 | 0.24 | 22 | 0.11 | 0.51 |
| g | Porphyromonas | 2.08E-02 | 0.24 | 21 | <u>0.67</u> | 0.49 |
| s | Prevotella | 2.70E-02 | 0.22 | 22 | <u>0.64</u> | 0.59 |
| g | Stomatobaculum | 3.59E-02 | 0.31 | 16 | <u>0.86</u> | 0.95 |
| g | Parvimonas | 3.88E-02 | 0.26 | 28 | <u>0.53</u> | 0.46 |

**Figure 4:**
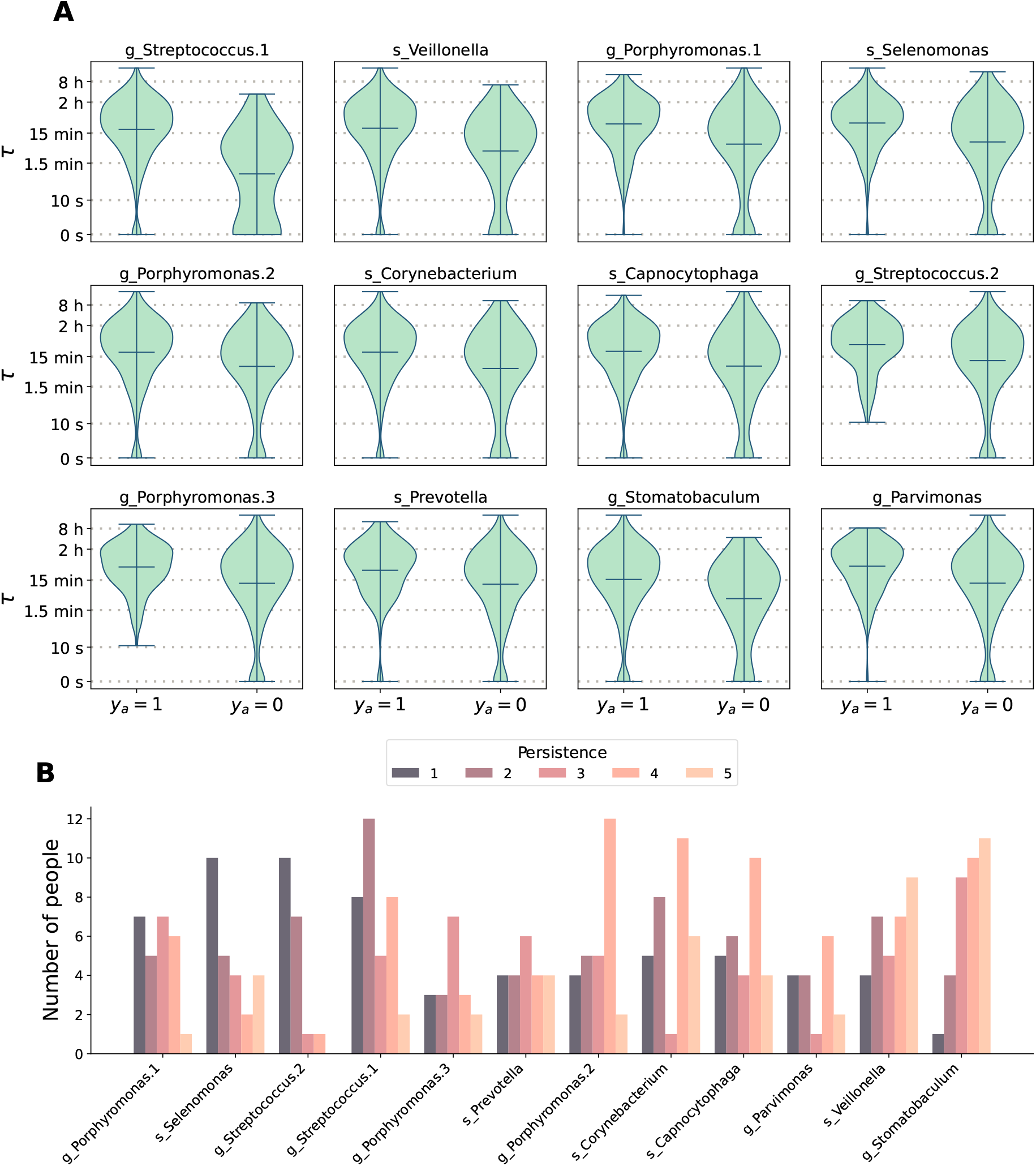
Taxa more strongly associated with the network structure. *Panel A*: Time spent in proximity by participants who share or do not share a given taxon. Each panel corresponds to one ASV and shows in logarithmic scale the histogram of time spent in proximity by participants who share (left-hand violin plot, *y*_*a*_ = 1) or do not share (right-hand violin plot, *y*_*a*_ = 0) a given taxon. The title reports the last taxonomic level available with “s” being the *species*, and “g” the *genus*. Note that we imposed log(0) = 0 to enhance readability. The plots show the two distributions for the 12 taxa for which the Mann-Whitney test is significant, after the Bonferroni correction. The notation *x*.1, *x*.2 is used to distinguish different taxa that share the same last available taxonomic level. *Panel B: Persistence of the taxa associated with the proximity network*. For each of the 12 significant variants shown in panel A, the bars show how many people were detected positive for that taxon in exactly 1 to 5 of their 5 samples (color-coded), *i*.*e*. how persistently each carrier hosted the variant rather than showing it only occasionally. ASVs are ordered by increasing global persistence from left to right.

### Association between network centrality and microbiota diversity

Previous studies showed that both in humans [8, 25] and animals [18], individuals with a larger social network have higher gut and oral microbiota diversity. We investigate this relation with our data, following Ref. [25]. Here, the authors introduce the *bacterial sharing network* by letting the weight of the edge (*ij*) be the number of shared taxa between *i* and *j* among the ones associated with the proximity network structure (those listed in Table 1, in our case). The authors then investigate the correlation between the eigenvector centralities in the proximity network and in the bacterial co-sharing network. Fig. 5 shows the results of this analysis on our data and displays a strong positive correlation (Spearman *r* = 0.52, with *p* value = 1.5 *·*10^*−*3^) between the centrality values obtained from the two networks. In the Supplementary information, we explore other centrality measures and show that the node strength in the proximity network *G*_p_ also correlates with the number of taxa displayed in Table 1 hosted by each node.

**Figure 5:**
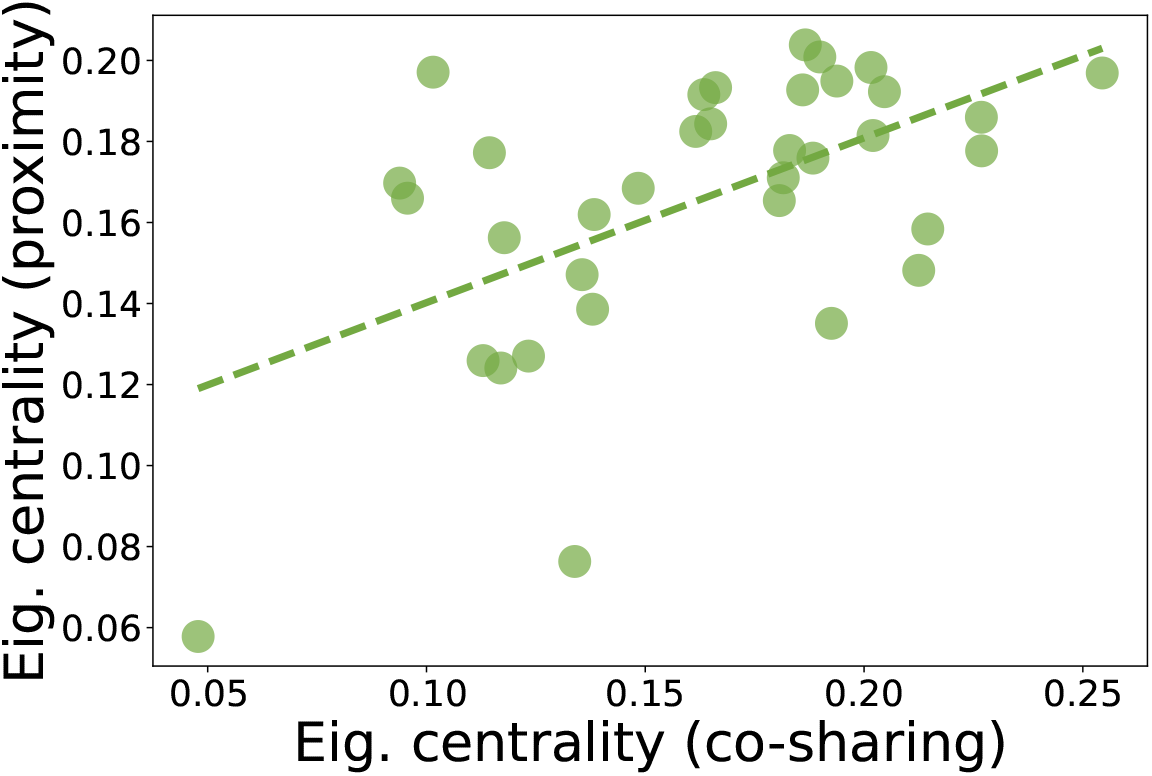
Association between node eigenvector centralities in the proximity and in the bacterial co-sharing networks. Each dot refers to an individual. The *y*-axis represents the node’s eigenvector centrality in the proximity network *G*_p_. The *x*-axis represents the node’s eigenvector centrality in the bacterial co-sharing network, whose edge weights are defined as the number of taxa from Table 1 shared by the edge nodes. The dashed green line is the result of the linear regression between the two centrality metrics. The Spearman correlation between the two variables is *r* = 0.52 with *p* value = 1.5·10^*−*3^.

## Discussion

Our study shows a statistically significant association between the composition of oral microbiota and close-range proximity of non-cohabiting children in a primary school. Our findings are qualitatively consistent with previous studies involving cohabiting individuals [22] and with recent, larger-scale studies focusing on gut or oral microbiome [27, 15] in isolated social settings, including studies using wearable proximity sensors in isolated communities [25].

The distinctive feature of our study is the choice of a highly non-isolated social environment – a primary school in a developed country – and the adoption of a comparatively low-resource approach. We use 16S rRNA gene sequencing and low-cost proximity sensors deployed for just a few weeks in the chosen social setting, without observing any of the contacts children have off-school with household members and their communities.

We remark that children were instructed to wear the proximity sensors also before and after school hours, including the weekend, but off-school proximity contributes negligibly to the measured contacts and was therefore excluded from our analysis. Of course, this might be due to lower compliance with sensor wearing outside of school hours, but it is important to remark that the proximity durations we observe are almost exclusively due to close-range proximity in the school.

We also show that information about the oral microbiota predicts sustained close-range proximity relations, with a prediction performance similar to that reported by other studies [26, 27] in isolated settings. This suggests that the oral microbiota structure is significantly shaped by close-range proximity, at least for some taxa. We also identify a set of ASVs whose presence in the oral microbiota correlates with prolonged close-range proximity. Possibly, these variants may be shared as a result of close-range social interactions at school—a hypothesis that our cross-sectional design cannot directly test, but that is consistent with the homophily observed in Fig. 2 and Table 1. – and we remark that several of these variants have also been associated with social interactions by other studies [25]. Moreover, we observe that the most central nodes in the proximity network have a higher number of these variants, in agreement with previous studies showing that individuals with a larger number of social interactions have a higher microbiota diversity [8, 25].

On the one hand, our results are consistent with the hypothesis that, at least for some taxa, sustained close-range proximity at school is associated with sharing of the oral microbiota, which is compatible with, though not proof of, social transmission. On the other hand, it is important to remark that time proximity at school, measured during our observation period, is probably a proxy for the time in proximity spent by the same individuals during the whole school year, as well as for other types of unobserved kinships among children. Since social influence and latent homophily on unobserved attributes – socio-economic attributes, dietary habits, etc. – generally cannot be teased apart, care needs to be taken in interpreting our results as evidence for the social transmission of oral microbiota, even assuming transmission is at play. In fact, it is important to stress that the proximity times we observe should not be regarded as causally relevant for transmission. For instance, the proximity times in Fig. 2 should not be interpreted as typical times required for microbiota transmission, as they only provide a definition of a social tie sufficiently strong to be associated, in our study, with the composition of the oral microbiota. Moreover, shared, non-social exposure – classroom air, canteen food, shared surfaces or drinking sources – could also drive co-acquisition of taxa among classmates without direct person-to-person contact, and our study is purely associative.

The time scales relevant for our study also need to be discussed. We observe that the microbiota of all individuals is essentially stationary over the 2 months spanning the days on which saliva samples were collected. This is compatible with two opposed scenarios: one in which the time scale of the presumed transmission is much longer than the duration of our study, and another in which this duration is much shorter. In the former case (long time scale needed for transmission), the duration of our study is insufficient for observing any significant microbiota dynamics determined by close-range proximity. In the latter case (short time scale needed for transmission), the microbiota of all individuals may have already achieved a stationary state, because our data collection started weeks after the beginning of the school year. In both cases, this study only observed a snapshot of a complex process that likely involves latent homophily on several unobserved attributes.

A further limitation is the resolution of 16S rRNA gene sequencing itself: it typically resolves taxa to the genus or species level (as reflected in Table 1, where several associations are only resolved to genus) and cannot distinguish between strains of the same species.

To overcome some of the limitations discussed above, future studies might envision a longer data collection period, with the goal of exposing the temporal dynamics of oral microbiota and identifying the time scales relevant for changes in microbiota determined by close-range proximity. Increasing the sampling frequency of oral microbiota might also be necessary to expose microbiota dynamics, and a more fine-grained shotgun metagenomic sequencing might be essential to overcome the limitations of 16S rRNA gene sequencing. Similarly, to deal with confounders and tease apart oral *microbiota association with close-range proximity*, such as the one observed in this study, and *microbiota transmission determined by close-range proximity*, future studies might enroll a larger cohort, possibly including the students’ households, and collect richer information about study participants, such as socio-economic attributes, dietary habits, and more.

## Methods

### Ethics

The Regional Committee for Medical and Health Research Ethics (REK East Norway) reviewed the research protocol and concluded it did not fall under the scope of “The Act on Medical and Health Research”, as the aim was not to directly generate knowledge about health and disease. Consequently, no REK approval was required for this study. The project received approval from the Data Protection Officer at Molde University College (September 2022). All re-identifiable data were stored at Molde University College and only de-identified data files were shared with research group members.

### Enrollment

Our study was conducted in a Norwegian primary school with 292 children, subdivided into 7 grades. Before recruitment commenced, the details of the study were discussed with the faculty at the primary school to ensure they had a full understanding of what the study entailed. Parents/guardians were invited to a meeting where the study was presented, and questions could be asked. Written information about the study was provided to the parents/guardians in two versions: one for the adults and another for the children, along with a consent form. Children were informed about the study orally in class. Inclusion required consent from both parents/guardians and assent from the child.

We invited 41 children aged 7-8 years (22 boys and 19 girls), 37 participated (21 boys and 16 girls). Due to sensor damage, however, the proximity data were collected for 35 children that constitute the cohort analyzed in this study. The experiment was conducted from September to November 2022. The children had a five-day school week with four to five class hours daily. During October 2022, the school had an autumn break with the duration of one week. After the data collection was completed, all children in the two classes were invited to an afternoon trip to the local cinema and a class trip to an aquarium in a neighboring city, both financed by the study. No distinction was made between children participating in the study and non-participating children.

### Experiment description

One dedicated researcher from our team served as the contact point for the children, their guardians, and the teachers. This researcher organized the collection of materials at the school throughout the whole data collection period, with visits to both classes every one to two weeks. Saliva samples were collected at 8 time points, with a one-week interval between the first five samples and a two-week interval between the last three. The samples were taken from September 12th to November 25th, 2022. Daily use of sensors overlapped this entire period. The proximity sensors had to be worn outside the clothing, making them visible to other children. They had a unique identifier to simplify their identification by each child. To avoid the possible stigmatization, children who did not participate were given the option to take saliva samples without storing them for analysis and to wear a sensor with the battery removed, i.e., without data recording. Four of the 41 children invited used this opportunity.

There were no compliance issues with either procedure. The children were highly motivated to give saliva samples and to use the sensors throughout the whole study period. Feedback from parents and teachers indicated that the children were self-motivated and independent. They also looked out for each other to ensure they remembered to wear the sensors, e.g., after gym when the sensors had to be taken off. The study did not impact the children’s routine, except for 10*−* 15 minutes set aside for collecting saliva samples on specific days. Our third-grade cohort participated in an after-school care program in the mornings (07 : 15 *−*08 : 20), and joined fourth-grade children in the program during the afternoons (13 : 00*−* 16 : 45). Most of the children in our cohort also live near each other and frequently spend time together outside of school hours.

### Saliva sampling

The children performed saliva sampling during class hours using the GeneFiX− Saliva Microbiome DNA collection kits (Isohelix, Cell Projects Ltd, Harrietsham, England)^1^ containing sample tubes with 1 ml stabilizing solution and funnel. They were required to not eat or drink during the 30 minutes before the procedure. All the steps in the saliva collection were performed simultaneously by all the children in each class, under the instruction of the dedicated researcher from our team. First, an envelope containing the collection kit and labeled with each child’s name was handed out. Next, the children followed the manufacturer’s instructions. Finally, the children placed the collection tube back into the envelope and delivered it to the researcher. The collection tube was premarked with a GeneFix− seven-digit serial number and a barcode, which were linked to each child’s study ID and the date of sampling. The saliva samples were stored at room temperature before DNA extraction. All samples were transported overnight from Molde University College to the laboratory at Oslo University Hospital HF in autumn 2022 and received in two batches on October 4th and November 30th.

### 16S rRNA gene sequencing and microbiota profiling

DNA from each saliva sample was extracted using the NAxtra− nucleic acid extraction kit (Lybe Scientific) following the manufacturer’s instructions, and then sent to the Norwegian Sequencing Center (Oslo, Norway) for 16S rRNA gene amplification, library preparation, and sequencing. In brief, the V3-V4 region of the 16S rRNA gene was amplified based on the two-step PCR procedure as described in the Illumina application note using primer sequences derived from Ref. [50]. Libraries were pooled and sequenced on an Illumina MiSeq platform (Illumina, San Diego, CA, USA) with 300 bp paired-end reads (v3 reagents). 20% PhiX control library was added to the libraries, and cluster density was reduced to 80% of regular levels. Base calling and production of demultiplexed FASTQ files were performed by running RTA v1.18.54.4 and bcl2fastq v2.20.0.422.

The demultiplexed FASTQ files were analyzed using QIIME2 version 2022.8 run via a SLURM script.

The analysis comprised the following steps:

1. Reads were demultiplexed and adapters trimmed using cutadapt with forward and reverse sequences CCTACGGGN-GGCWGCAG and GACTACHVGGGTATCTAATCC. To remove low-quality calls at the ends of the reads, adapter-trimmed reads were truncated at 250 bp for the forward read and 190 bp for the reverse read.
2. Denoising and amplicon sequence variant (ASV) inference was carried out using DADA2.
3. Taxonomic classification of the ASVs was carried out using the feature-classifier classify-sklearn command with SILVA release 138 as the reference database.
4. A phylogenetic tree was estimated using the align-to-tree-mafft-fasttree command, which generates a MAFFT alignment from reads, and fasttree was used to estimate a phylogenetic tree from these alignments.
5. To remove possible contaminations (taxa present in the negative control water sample), a vector was created with all ASVs except the ones present in the negative control. This vector was then applied to the ‘prune taxa()’ function from the R phyloseq package on the ‘phyloseq’ object.

The QIIME2 analysis SLURM script is available on GitHub at github.com/CBGOUS/.

### Proximity network construction

The proximity network was measured using the `SocioPatterns` wireless proximity sensors [30, 31]. These are wearable, unobtrusive devices embedded in a badge worn by the participants using a lanyard. Each sensor emits low-power radio packets at known power on a radio channel within the ISM radio band at 2.4 GHz, at a rate of approximately 40 packets per second. When two sensors exchange at least one information packet, the receiving sensor records the unique identifier of the sender, the interaction time-stamp, and the signal attenuation value. Proximity within approximately 1.5 meters is obtained by filtering the measurements according to their attenuation. Moreover, each sensor periodically records some status properties that are used during data cleaning to determine the correct functioning of the sensor. Among these, an accelerometer detects whether the sensor is moving or not. This signal can be used to filter out contact data collected while the sensor is not moving which is likely because it was not worn. During the measurement period, the sensors remained active and continuously recorded proximity data. The field team reported a high level of compliance throughout the experiment by all children, corroborated by the readings on the onboserd sensor’s accelerometer. This technology has been extensively tested and used for quantifying human contacts in a variety of contexts, including schools [39, 41], hospitals [44] and African rural villages [48, 47], among others.

Based on a preliminary analysis, we observed a rapid deterioration in the data quality collected by the proximity sensors over successive waves, primarily due to damage. Consequently, we opted to only utilize the initial two measurements as reliable inputs for our analysis. These two measurements show a high correlation in the contact patterns: the Spearman correlation between the weights is *r* = 0.25 with *p* value 7 *·*10^*−*14^. These correlations, paired with the high compliance observed in these measurements, make them reliable data sources. Although children were instructed to wear the sensors before and after school hours as well, contacts outside school hours were excluded upstream of the analysis presented here, so all reported proximity durations reflect school-hours contact only.

## Acknowledgments

We thank the children for their participation in the study, the parents for their invaluable help ensuring that the sensors were worn every day, the rector and teachers for their support and practical assistance in executing the study.

## Funding

This project was funded by the Norwegian Institute of Public Health and the Molde University College. LD, CC and DP acknowledge the support from the Lagrange project of Fondazione CRT. LD and CC acknowledge the support from Fondation Botnar (EPFL COVID-19 Real Time Epidemiology I-DAIR Pathfinder).

## Author contributions

CSN and CC designed the study. SMW and ASF supervised the data collection in the field. XB, SR, and AM carried out the microbiota profiling. CC, LD, and DP performed the data analysis. LD, CC, and XB wrote the first version of the manuscript. All authors interpreted the results, contributed to the final version of the manuscript, and approved the final version of the manuscript.

## Supplementary information

### S.1 Robustness to the choice taxon-filtering thresholds

We investigate the robustness of our results by filtering rare and low persistent variants. The first filter is on *prevalence*, which drops variants observed in at most *p* ∈ {0, 1, 10} samples across the entire population. For *p* = 0 there is no filtering, *p* = 1 singletons are removed, whereas *p* = 10 is a more stringent filter on rare variants. The second filter is on *persistence* and it sets to zero, independently for each individual, the abundance of variants recurring in fewer than *q* ∈ {1, 3} samples. For *q* = 1 no filtering is applied, while for *q* = 3 only variants detected in the majority of samples are retained.

For each of the four candidate distance metrics (Jaccard, Bray-Curtis, weighted UniFrac, and Unweighted UniFrac), we computed the pairwise sample-distance matrix at every filtering level and report the minimum Spearman correlation between any pair of filtering levels: 0.87 for Jaccard, 0.97 for Bray-Curtis, 0.95 for weighted UniFrac, and 0.42 for unweighted UniFrac. Because unweighted UniFrac is markedly less robust to the choice of filtering thresholds than the other three metrics, we exclude it from the robustness analysis.

Figure S.1 reproduces the main-text result of Fig. 1A, showing that microbiota samples collected from the same individual at different times are more similar than samples collected from different individuals. The figure shows that this result is robust to both persistence and prevalence thresholds and the separation between the two distributions actually increases with both threshold values. Figure S.2 shows that the main-text result of Fig. 2 – lower microbiota distance among pairs with longer contact duration, relative to the null model – holds across all six filtering levels, for each of the three retained distance metrics.

Finally, Table S.1 reports the PERMANOVA results for all three distance metrics and all six filtering levels, grouping samples by individual identity and by sampling day. Grouping by individual identity is significant for every metric and filtering level, while grouping by sampling day is not significant for Jaccard or Bray-Curtis at any filtering level; for weighted UniFrac is significant, but with a substantially smaller effect size than individual identity, confirming the robustness of the results shown in the main text.

**Figure S.1:**
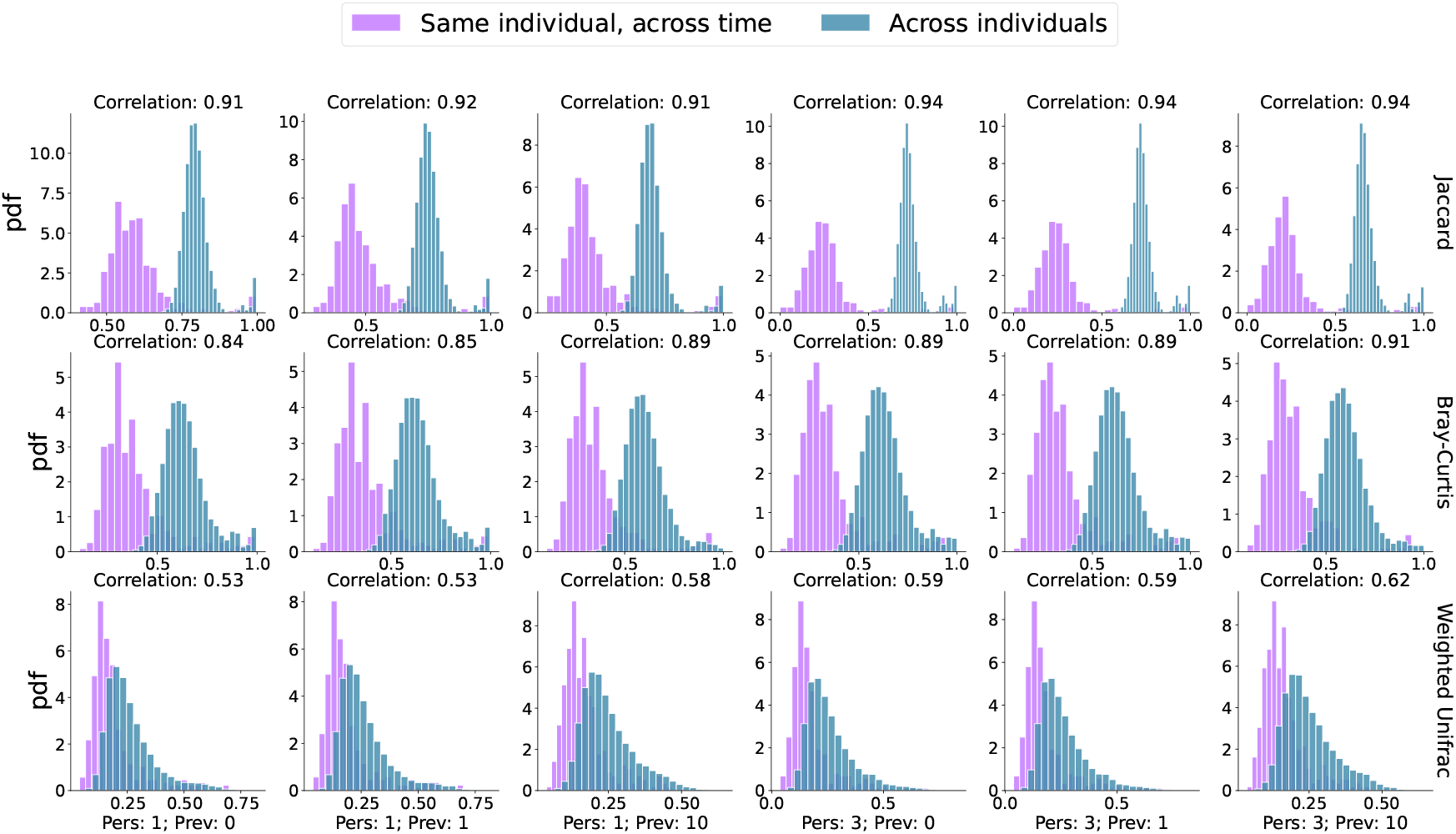
Robustness of the similarity of microbiota across individuals and across time for the same individual to the prevalence and persistence thresholds. Histograms of microbiota distance across individuals and across time for the same individual, for three distance metrics and for the six combinations of persistence and prevalence values. For each frame, the histograms represent the distance between all measurements involving the same person (“Same individual, across time”, purple) and between all measurements involving different individuals (“Across individuals”, blue). The plots in the same column correspond to the same persistence and prevalence thresholds. The title of each panel reports the rank-biserial correlation of the Mann-Whitney test.

**Figure S.2:**
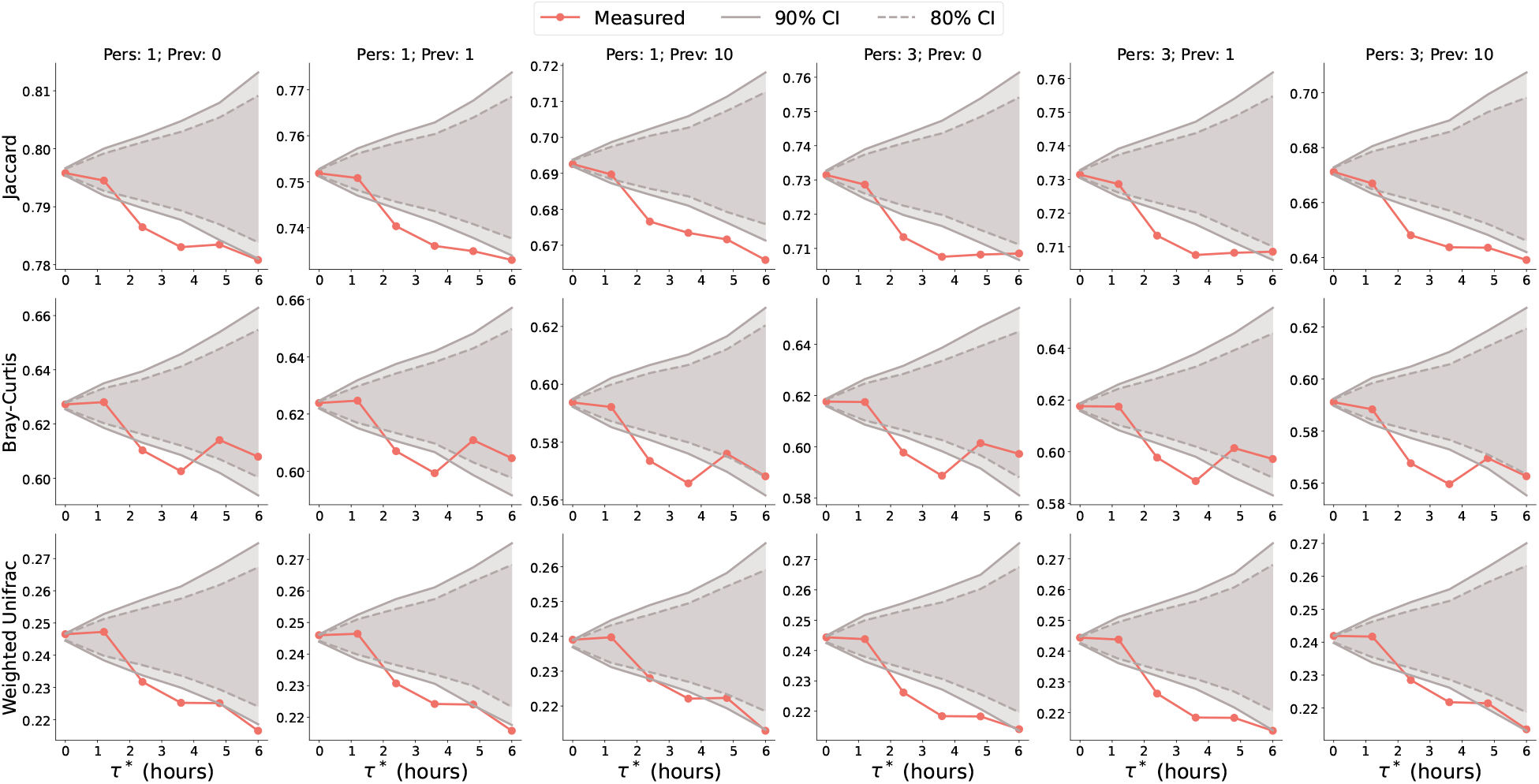
Robustness of the strong-tie analysis across distance metrics and filtering levels. Strong ties have a duration *τ* larger than the threshold *τ*\*, spanning the *x*-axis. The *y*-axis shows the average microbiota distance (weight of *G*_m_) between nodes forming strong edges, for the corresponding metric. The two style-coded gray lines are the 80% and 90% confidence intervals obtained over 1500 realizations of the null model randomizing the association between the weights of *G*_p_ and *G*_m_. Proximity strength increases from left to right, whereas microbiota similarity increases from top to bottom. The plots in the same column correspond to the same persistence and prevalence thresholds.

**Table S1:** PERMANOVA results (*R*^2^ and *p*-value) across distance metrics and filtering levels.

| Distance | Level | Same individual, across time |  | Across individuals |  |
| --- | --- | --- | --- | --- | --- |
| | | $R^2$ | $p$ | $R^2$ | $p$ |
| Jaccard | Pers. 1, Th. 0 | 0.018 | 1.000 | 0.547 | 0.001 |
|  | Pers. 1, Th. 1 | 0.017 | 1.000 | 0.650 | 0.001 |
|  | Pers. 1, Th. 10 | 0.018 | 0.999 | 0.670 | 0.001 |
|  | Pers. 3, Th. 0 | 0.007 | 1.000 | 0.874 | 0.001 |
|  | Pers. 3, Th. 1 | 0.007 | 1.000 | 0.874 | 0.001 |
|  | Pers. 3, Th. 10 | 0.007 | 1.000 | 0.869 | 0.001 |
| Bray-Curtis | Pers. 1, Th. 0 | 0.025 | 0.220 | 0.689 | 0.001 |
|  | Pers. 1, Th. 1 | 0.025 | 0.203 | 0.695 | 0.001 |
|  | Pers. 1, Th. 10 | 0.029 | 0.069 | 0.710 | 0.001 |
|  | Pers. 3, Th. 0 | 0.025 | 0.267 | 0.741 | 0.001 |
|  | Pers. 3, Th. 1 | 0.025 | 0.276 | 0.741 | 0.001 |
|  | Pers. 3, Th. 10 | 0.028 | 0.094 | 0.743 | 0.001 |
| weighted UniFrac | Pers. 1, Th. 0 | 0.041 | 0.021 | 0.549 | 0.001 |
|  | Pers. 1, Th. 1 | 0.041 | 0.011 | 0.553 | 0.001 |
|  | Pers. 1, Th. 10 | 0.042 | 0.013 | 0.607 | 0.001 |
|  | Pers. 3, Th. 0 | 0.043 | 0.019 | 0.608 | 0.001 |
|  | Pers. 3, Th. 1 | 0.043 | 0.007 | 0.608 | 0.001 |
|  | Pers. 3, Th. 10 | 0.041 | 0.024 | 0.644 | 0.001 |

### S.2 Inference of the proximity network from the microbiota with a random forest classifier

We follow the procedure described in **Inference of the proximity network from the microbiota** and use a random forest classifier instead of a support vector machine to test the robustness of our result. The test is conducted with the same parameters, and it only differs in the use of the classifier. Figure S.3 shows the ROC curve obtained for 100 random splits of the train and test sets. The average AUC equals 0.74.

**Figure S.3:**
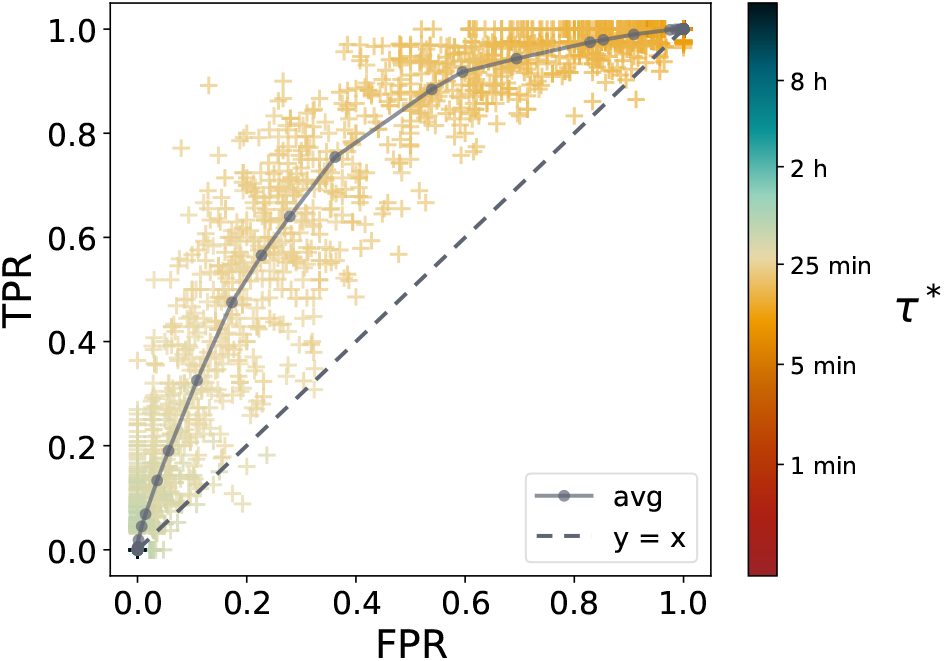
ROC curve for the random forest binary classifier used to infer the proximity links based on microbiota profiles. We follow the procedure detailed for Figure 3 using a random forest classifier to predict whether the weight (duration in proximity) of a given edge (*ij*) exceeds a threshold *τ*\*. The results are obtained for 100 random split of the training/test sets, for each of the 250 logarithmically spaced values of *τ*\* we scan. Every cross in the plot corresponds to one realization of the training and test sets. The *x* coordinate is the false positive rate (FPR) and the *y* coordinate is the true positive rate (TPR). Markers are color-coded according to the value of the threshold *τ*\*. The solid gray line represents the average over the random splits and has an area under the curve auc = 0.74. The dashed line is the curve *y* = *x* for a random-guess binary classifier.

### S.3 Analysis of microbial alpha diversity

Figure S.4 shows the microbial alpha diversity in the five measurements analyzed in the main text, at every filtering level described above. Using different metrics (number of observed ASVs, Shannon, Fisher, and Simpson diversity indices), we show that the alpha microbial diversity remains stable over the five measurements. No pair of measurements has highly significant (Mann-Whitney *p <* 0.05 with Bonferroni correction) differences in the distributions, according to any diversity metric, at any filtering level.

### S.4 Analysis of temporal turnover of shared variants

For every pair of children and every pair of sampling days, we computed the Jaccard similarity between the sets of variants shared by the pair on those two days, grouping the results by the number of days elapsed between sampling events (Fig. S.5). The median similarity remains stable across all day gaps considered (4 to 25 days), fluctuating only between 0.54 and 0.60, with no monotonic trend, indicating that there is no systematic temporal turnover in the sets of shared variants.

### S.5 Correlation between proximity and bacterial sharing network

We investigate the correlation between the social network *G*_p_, measured using the proximity sensors, and the bacterial sharing network. We recall that, for this network, the weights of an edge (*ij*) are defined as the number of variants appearing in Table 1 shared by the nodes *i* and *j*. Figure S.6A shows that the edge weights of the two networks are highly correlated (Spearman *r* = 0.44 and *p* value below machine precision). The subsequent panels complement Figure 5 by considering alternative node centrality measures for which we obtain strong across the two networks: random walk betweenness (Spearman *r* = 0.51, *p* value = 1.4·10^*−*3^),closeness (Spearman *r* = 0.51, *p* value = 1.7·10^*−*3^), and strength (Spearman *r* = 0.51, *p* value = 1.7·10^*−*3^).

**Figure S.4:**
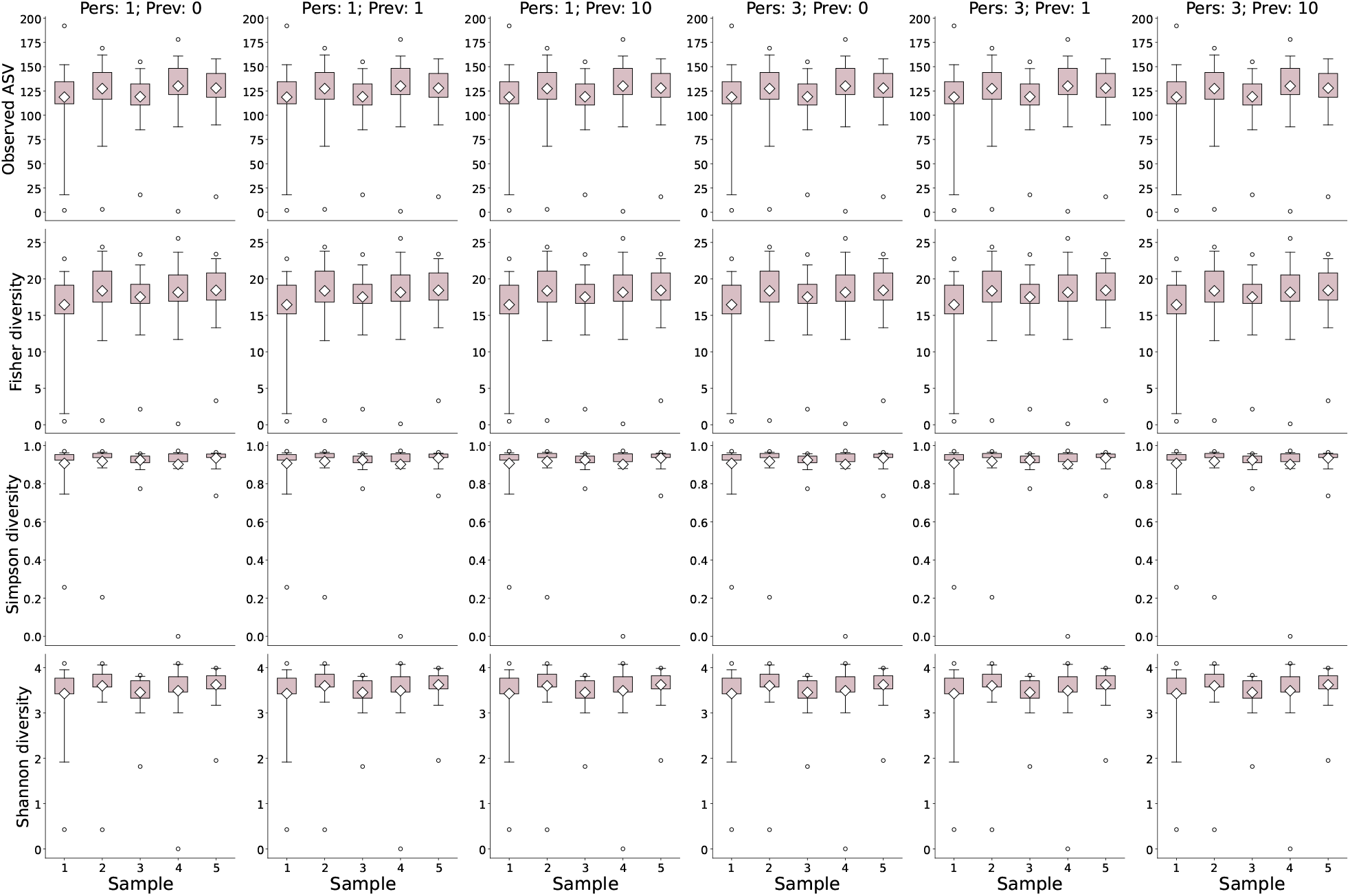
Alpha microbial diversity. Boxplot of the alpha diversity according to four different diversity measures indicated in the frames’ titles, for the six taxon-filtering levels considered (columns). The *x*-axis spans the five measurements considered in the main text. The whiskers indicate the 1 *−* 99 confidence interval, while the white diamond is the sample average.

**Figure S.5:**
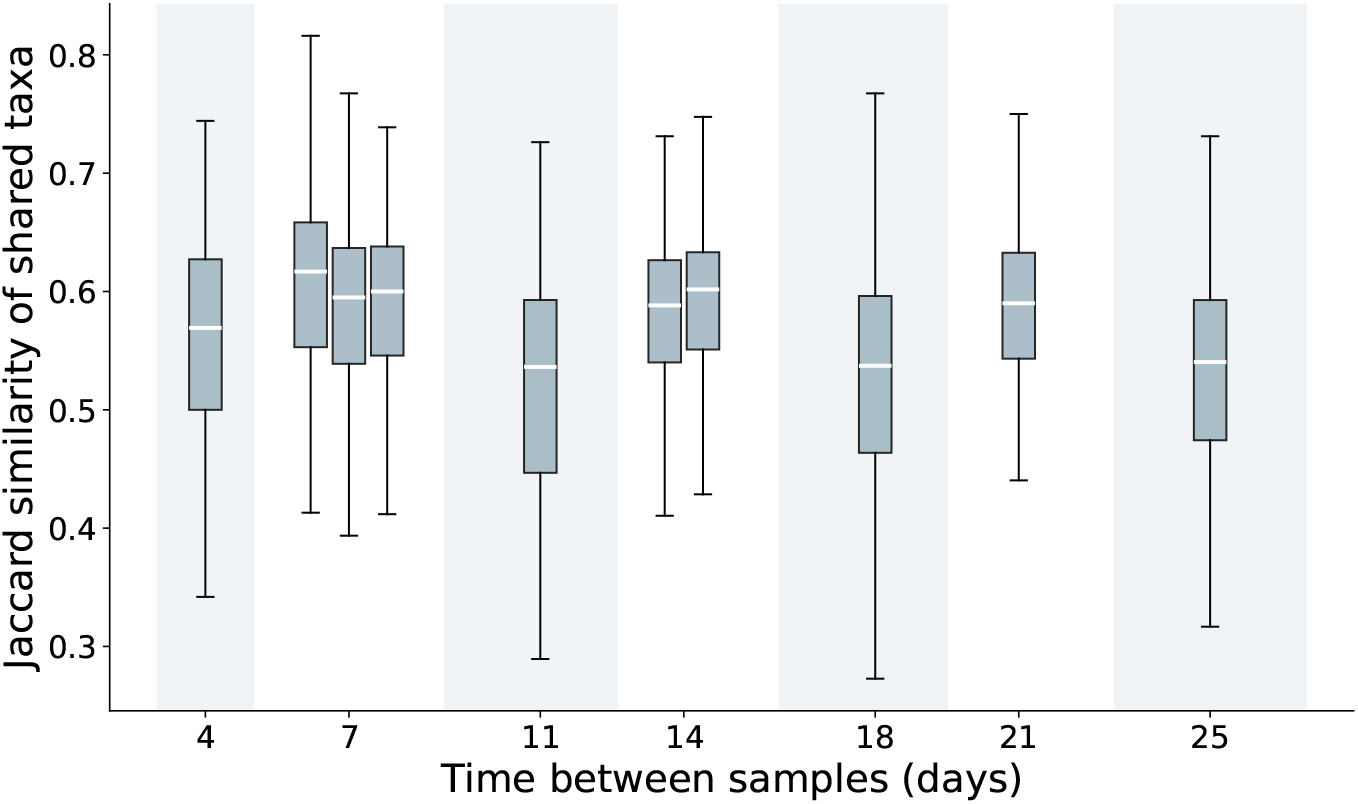
Temporal turnover of shared variants. For every pair of children, we computed the Jaccard similarity between the sets of variants shared on two sampling days and grouped the results by the number of days elapsed between the two sampling events.

**Figure S.6:**
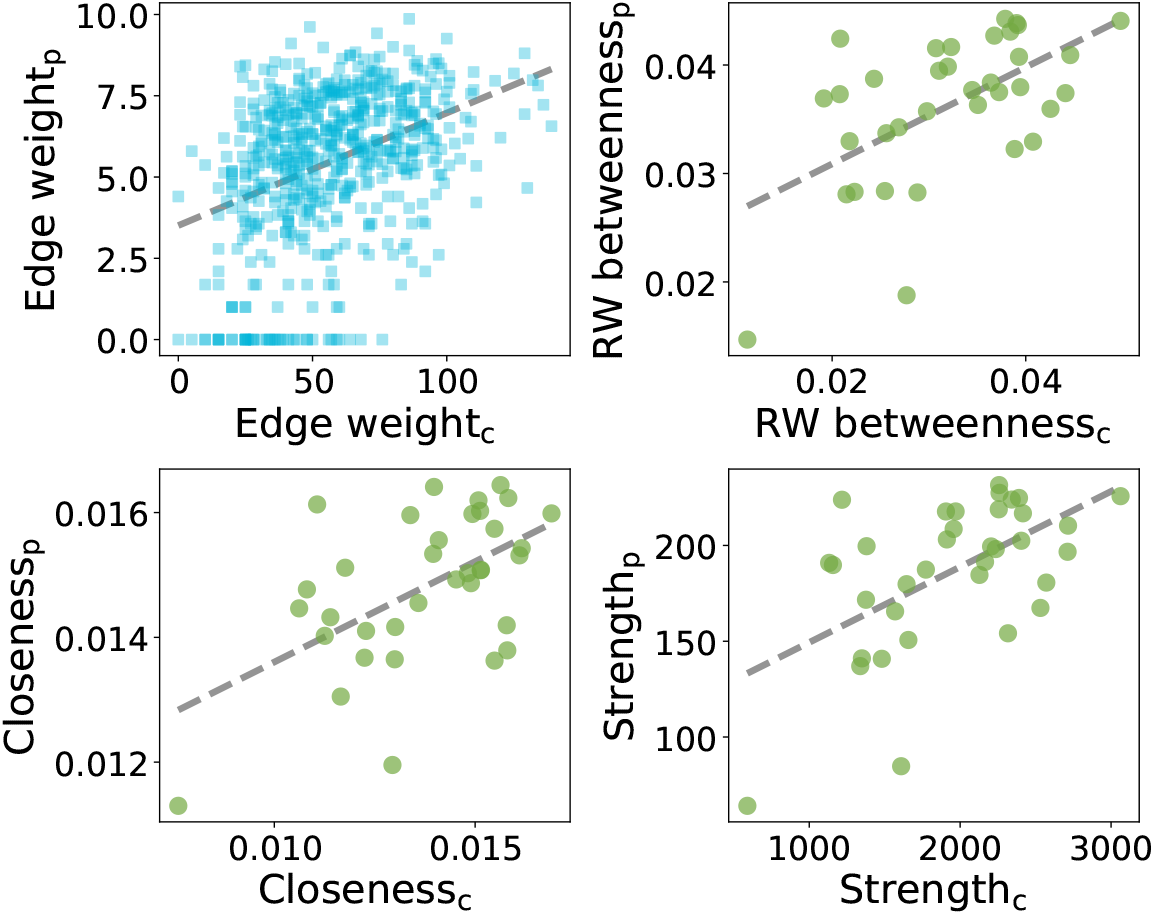
Correlation between the proximity network and the bacterial sharing networks. *Panel A*: correlation between the edge weights in the two networks. *Panels B, C, D*: correlation of different node centrality measures (random walk betweenness, closeness, and strength) across the two networks. The subscript “c” stands for *co-sharing*, while the subscript “p” for *proximity*.

## Footnotes

1 www.isolhelix.com

## Notes

https://github.com/lorenzodallamico/MicrobiotaSocioPatterns

### Competing Interest Statement

The authors have declared no competing interest.

### Summary of Updates

This version of the manuscript handled a more detailed robustness analysis on the cleaning of microbial data as well as on the dissimilarity metrics adopted. Some parts were extensively rewritten to improve readability.

